# neoantigenR: An annotation based pipeline for tumor neoantigen identification from sequencing data

**DOI:** 10.1101/171843

**Authors:** Shaojun Tang, Subha Madhavan

## Abstract

Studies indicate that more than 90% of human genes are alternatively spliced, suggesting the complexity of the transcriptome assembly and analysis. The splicing process is often disrupted, resulting in both functional and non-functional end-products (Sveen et al. 2016) in many cancers. Harnessing the immune system to fight against malignant cancers carrying aberrantly mutated or spliced products is becoming a promising approach to cancer therapy. Advances in immune checkpoint blockade have elicited adaptive immune responses with promising clinical responses to treatments against human malignancies (Tumor Neoantigens in Personalized Cancer Immunotherapy 2017). Emerging data suggest that recognition of patient-specific mutation-associated cancer antigens (i.e. from alternative splicing isoforms) may allow scientists to dissect the immune response in the activity of clinical immunotherapies (Schumacher and Schreiber 2015). The advent of high-throughput sequencing technology has provided a comprehensive view of both splicing aberrations and somatic mutations across a range of human malignancies, allowing for a deeper understanding of the interplay of various disease mechanisms.

Meanwhile, studies show that the number of transcript isoforms reported to date may be limited by the short-read sequencing due to the inherit limitation of transcriptome reconstruction algorithms, whereas long-read sequencing is able to significantly improve the detection of alternative splicing variants since there is no need to assemble full-length transcripts from short reads. The analysis of these high-throughput long-read sequencing data may permit a systematic view of tumor specific peptide epitopes (also known as neoantigens) that could serve as targets for immunotherapy (Tumor Neoantigens in Personalized Cancer Immunotherapy 2017).

Currently, there is no software pipeline available that can efficiently produce mutation-associated cancer antigens from raw high-throughput sequencing data on patient tumor DNA (The Problem with Neoantigen Prediction 2017). In addressing this issue, we introduce a R package that allows the discoveries of peptide epitope candidates, which are the tumor-specific peptide fragments containing potential functional neoantigens. These peptide epitopes consist of structure variants including insertion, deletions, alternative sequences, and peptides from nonsynonymous mutations. Analysis of these precursor candidates with widely used tools such as netMHC allows for the accurate in-silico prediction of neoantigens. The pipeline named neoantigeR is currently hosted in https://github.com/ICBI/neoantigeR.

## Background

Alternative splicing shapes the mammalian transcriptomes as many RNA molecules undergo multiple alternative splicing events. This process is crucial in building transcriptome complexity and proteome diversity (Tazi, Bakkour, and Stamm 2009). Since long transcripts are often spliced into multiple alternative splicing variants, identification of transcript isoform and quantification of transcript-level expression from RNA-Seq data is a considerably difficult problem (Tang et al. 2016). It is usually unclear which reads are included in the particular exon of which transcript, and it’s hard to decipher the true complexity of the transcriptome. High-throughput RNA sequencing (RNA-Seq) enables characterization and quantification of transcriptome as well as detection of patterns from alternative splicing (Tilgner et al. 2015; Treutlein et al. 2014).

Though ongoing advances are rapidly yielding increasing read lengths, a technical hurdle remains in identifying the degree to which differences in read length influence various transcriptome analyses. The long-read sequencing approach overcomes these limitations by sequencing thousands of independent full-length mRNA transcripts. This technology also has many vital benefits, including lower mapping bias and reduced ambiguity in assigning reads to genomic elements, e.g. mRNA transcript (Cho et al. 2014; “Assessment of Highly Complex Alternative Splicing of Neurexins Performed with SMRT Sequencing” 2014). Detecting such splicing events could dramatically improve our understanding of the alternative splicing which is often associated with cancer (Cho et al. 2014). Furthermore, the identification of alternative splicing, fusion events, and novel transcripts will bring us increasingly closer to direct measurement of these events. In fact, long-read sequencing has demonstrated the capability to accurately capture full-length transcripts in an annotation-free manner and identified novel splicing patterns not reported based on short reads.

Ribosomal profiling indicates that many regions of transcriptome such as 5’ UTR and lincRNA - which were previously thought to be noncoding - are actually also translated. The discovery of pervasive translation of non-canonical transcripts is hampered by the short-read sequencing since popular short-read transcript assembly algorithms were mostly based on canonical transcripts. Consequently, long-read sequencing produces full-length transcripts without reliance on transcript assembly algorithm (which would greatly accelerate the detection of these non-canonical products). Current studies suggest that long-read sequencing is capable of recovering repetitive sequences, indentations, non-canonical transcripts from intergenic or intragenic regions that are lacking in short-read sequencing. The application of long-read sequencing will thus substantially facilitate the discovery of novel protein isoforms produced in experimental conditions.

Cancer immunotherapy has gained widespread popularity as shown by recent clinical application of checkpoint blockade inhibition. To date, multiple biomarkers have been associated with improved clinical outcomes in cancer patients receiving anti-PD-1 and anti-CTLA-4 therapy. Most of these biomarkers have characterized the immune status of the tumor microenvironment. Tumor mutational burden may serve as an additional, non-redundant biomarker for prediction of clinical benefit to both immunotherapy and targeted therapy. High mutational burden correlates to a greater number of neoantigens and tumor recognition by the innate and adaptive immune system, which is associated with clinical benefit from immune checkpoint therapy. Intriguingly, previous studies demonstrate that somatic mutations are strongly associated with diverse alternative splicing events (Furney et al. 2013), including alternative terminal exon usage, intron retention, and cryptic splicing within exons of both protein coding and noncoding genes.

Available data support the potential for predicting the clinical benefit of immunotherapy and targeted therapy based on molecular profiles in the baseline tumor microenvironment. Anti-tumor response and survival with checkpoint inhibitor immunotherapy has been associated with a series of likely interrelated factors including tumor PD-L1 expression, increased CD8 T-cell infiltration, interferon-gamma gene expression signature and high tumor mutational burden. Immunotherapies enable the endogenous T cells to recognize and destroy cancer cells. Many studies have demonstrated the efficiency of the therapy in a wide range of human malignancies (Schumacher and Schreiber 2015). The greater number of somatic mutations in human tumors varies among different patients, and some of these somatic mutations, if occurred in the coding region, might disrupt the opening reading frame of the translated protein. Therefore, it is believed that the resulting patient-specific peptide sequences (neoantigens) contribute to the clinical benefits of the immunotherapies (Schumacher and Schreiber 2015).

This study results in an innovative software package to facilitate the identification of disease/sample specific protein isoforms using both short read or long read high-throughput sequencing data and an existing protein database. By expanding the focus from somatic point mutations using pipelines such as pVAC-Seq (the only tool available to generate epitope candidates to date) to the entire neoantigen landscape, we aim to uncover important splicing aberrance and genomic abnormities which may cause larger changes in epitope binding affinity.

In addition, our epitope precursor candidates could be used as input for existing tools to allow precise binding affinity prediction. A number of software tools such as (IEDB) (Beaver, Bourne, and Ponomarenko 2007), EpiBot (Duarte et al. 2015), and EpiToolKit (Feldhahn et al. 2008) compile the results from individual epitope prediction algorithms to improve the prediction accuracy.

## Methods

The neoantigen prediction pipeline requires a very widely used standard data input from next-generation sequencing assays. In the simplest nontrivial scenario, the pipeline only needs a gene prediction output file in GFF format (Fig. 1). NeoantigenR will use the input GFF file to extract open reading frames (ORFs) from the predicted isoforms and compare them with reference protein database, typically from Uniprot Swiss-Prot or a previously utilized customized protein database. If the sequencing data is from long read sequencing, the input files will contain the mandatory gene prediction file (GFF format) and an optional raw sequence read file (FASTA format).

**Fig. 1.**
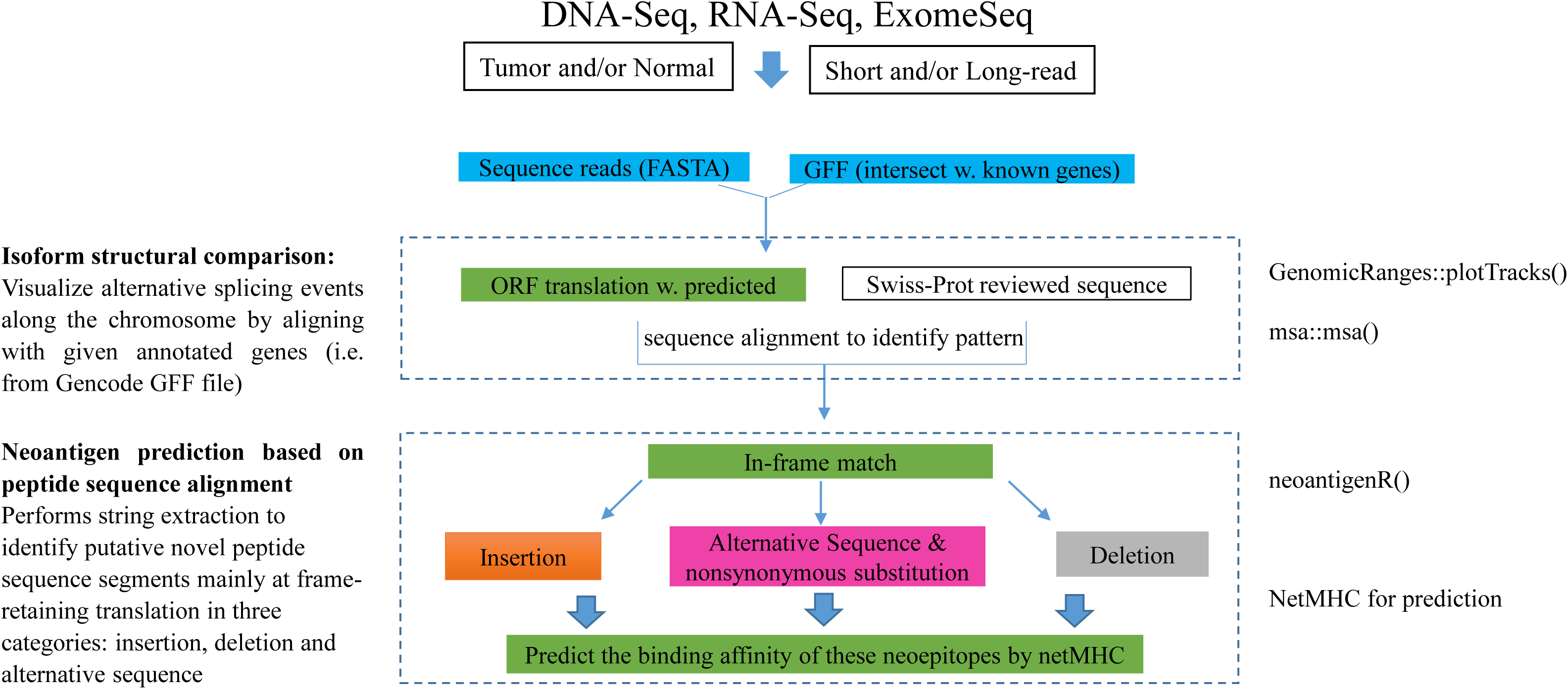
Flowchart of the discovery of novel peptides from alternative splicing in neoantigeR: This figures illustrates the methodology for neoantigenR pipeline construction. After proper preparation of input files, the predicted transcripts are translated and compared with canonical protein sequence for the identification of alternatively spliced peptides as the neoantigen candidates.

NeoantigenR implements individual modules via: sequence alignment and isoform calling, epitope prediction, and neo-antigen candidate scoring using existing tools such as netMHC (Nielsen and Andreatta 2016) (Fig. 1). The following paragraphs describe our unique analysis methodology from preparation of input files for the three modules of the pipeline.

### Preparing input data: reference genome alignment, gene-calling and annotation

As described above, neoantigenR relies on gene prediction input generated from the analysis of high-throughput parallel sequencing data including short-read RNA-Seq or long-read PacBio SMRT sequencing data. Generally, these data can be easily obtained from existing aligning and gene calling tools. Here, we outline example preparatory steps to generate these input data.

Reference genome sequence alignment was performed using the Bowtie2 (Langmead and Salzberg 2012) for aligning of original raw sequences (FASTQ files) to obtain SAM/BAM files. In brief, the Bowtie tool (release 2.1) was used for alignment with default parameters. The resulting alignments (in BAM format) file was subsequently used as the input to the gene calling tool Cufflinks (Trapnell et al. 2010) with de-novo gene-finding mode (no gene annotation is provided) and default parameters. Cufflinks accepted aligned RNA-Seq reads (SAM/BAM format) and assembled the alignments into a parsimonious set of transcripts. A gene/transcript prediction file in GFF format was then produced.

After sequence alignments and gene calling, annotation was performed by matching the transcript/gene prediction file (GFF) with an existing gene prediction (i.e. Gencode human release v19) using the following method: BEDTools (utilizing the ‘intersect’ utility function) was run on the above two files with parameter ‘-wo’ and default setting for the other parameters (Quinlan and Hall 2010). This command produced a gene-calling output file (Cufflink prediction output) and an existing gene annotation file (both in GFF file) and outputted a result file containing the intersection of the two files. The resulting file contained important information about the overlap of an exonic region between a predicted isoform comparing to a known gene.

### Isoform structural comparison to visualize alternative splicing events

This module aims to provide a structured visualization framework to plot gene/isoforms along genomic coordinates. It also allows for the integration of publicly available genomic annotation data from sources like UCSC. Using the predicted and given gene annotation, the pipelines pass the start and end positions of each annotation feature in the track, and also supply the exon, transcript, and gene identifiers for each item to create the transcript groupings (“Visualizing Genomic Data Using Gviz and Bioconductor - Springer” 2017). Such a visualization can be particularly useful for detecting alternative splicing events by comparing the predicted isoforms vs. annotated genes (i.e. from Gencode human v19 annotations) along a chromosome.

### Neoantigen prediction based on peptide sequence alignment

To streamline the identification of neoantigens, we will first build a protein sequence FASTA file that contains six ORFs of all predicted isoform. The protein sequence could be translated from a given reference genome or from original provided sequence reads. For instance, if the PacBio long-read sequencing full-length sequence reads are used, a direct in-silico translation will be performed on the input single molecule, real-time (SMRT) sequences (Kim et al. 2014). Meanwhile, peptide sequences for matched known genes are retrieved from Uniprot Swiss-Prot (Wu et al. 2006). Multiple protein sequence alignment will be performed using R msa package (Bodenhofer et al. 2015) based on commonly used methods such as ClustalW, T-Coffee and MUSCLE (Edgar 2004). After the protein sequence alignment, the pipeline will parse the alignment output and produce alternative peptide sequences that will be used as the basis for neoantigen prediction.

To begin with, the pipeline neoantigenR requires a continuous exact match of a certain number of amino acids on each side of the alternatively spliced sequence (the left and right anchor sequences respective, see Fig. 2), or anchor sequence in this study. It is straightforward to detect a sequence fragment absence (deletion) or a newly added sequence fragment (insertion) in the given predicted protein isoform by comparing it to the matched reference protein. Otherwise, sequences are translated from alternative exons, introns, splice sites or from same genomic locations with nonsynonymous mutations. The choice of an appropriate length of anchor sequence is crucial for the accuracy of detections. If the anchor sequence is too short, we may encounter a large number of falsely predicted alternative spliced regions and truly spliced regions may be missed. Therefore, we have conducted a statistical analysis to derive a recommended anchor sequence size. Firstly, we compute the probability of obtaining a match between one or more peptides of length SP amino acids from a query protein of length L random amino acids and a database of length D (DP random peptides of length SP was calculated as follows) (Silvanovich et al. 2006).

**Fig. 2.**
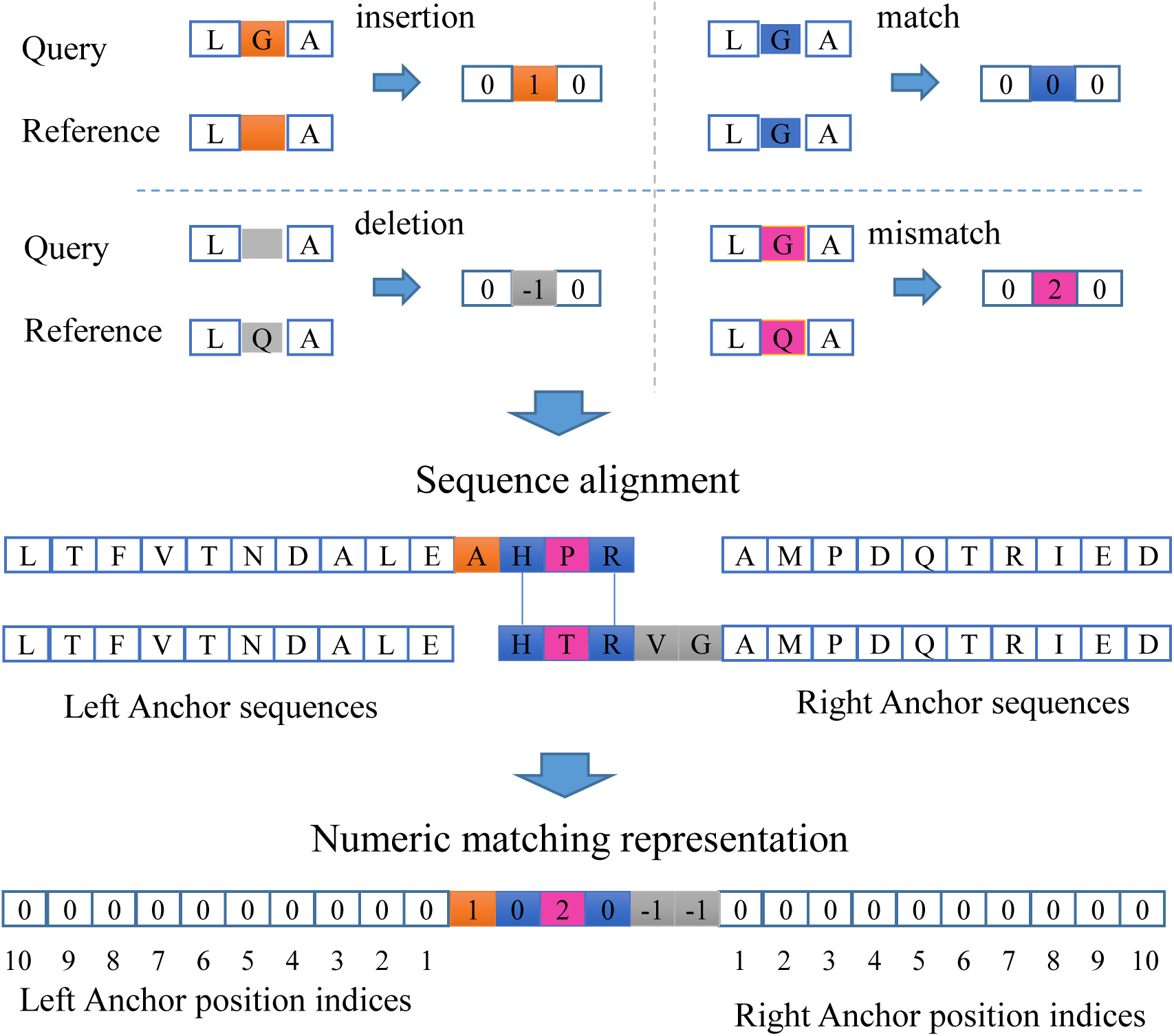
The protein sequencing alignment algorithm: sequences translated from predicted genes are aligned with reference protein database or protein sequence from matched normal samples. Protein sequences containing mismatched peptide sequence, deletions and insertions are identified assuming that there are continuous exact matches in both flanking regions.

NQ = 20^SP^, the number of possible unique short peptides of length SP.

ND = D– SP + 1, the number of short peptides in a database protein.

The probability of an arbitrary peptide of length SP is same as query sequence is: 1/20^SP^

The probability of an arbitrary peptide of length SP is different from query sequence is: (20^SP^-1)/20^SP^

The probability of a given short peptide is not in the database sequences = The probability of drawing ND peptides of length SP from database sequence with no match: DP=[(20^SP^-1) / 20^SP^]^ND^ the probability of a given short peptide occurring at least once in the database sequences QP = 1 – DP= 1-[(20^SP^-1) / 20^SP^]^ND^

Based on the above scenarios, the chance of observing a given 10 amino acid sequence peptides at least once in a human protein sequence (average protein sequence length is 480 amino acids) is approximately QP = 1 – DP= 1-[(20^10^-1) / 20^10^]^471^ ~ 0, even if we search a 10 amino acid sequence in the entire human protein database (480x20,000)=1–DP=1-[(20^10^-1)/20^10^]^9600000^<1-(0.999999999)^9600000^<1-0.99=0.01. Therefore, the choice of an anchor size of 10 amino acids is sufficient to maintain the uniqueness of sequence match.

After choosing the proper anchor sequence length, neoantigenR performs string extraction to identify putative novel peptide sequence segments mainly at frame-retaining translation in three categories: insertion, deletion and alternative sequence (Fig. 2.) using the following steps: (1) for each input peptide sequence, a vector or an object from the AAStringSet aligns it with each of the annotated isoform sequence of the matching protein; (2) the alignment is performed by R function ‘msa’ with default parameters; (3) the alignment result is converted to a string of alignment identity for each amino acid using R function msaConvert; (4) a final alignment matrix is composed by translating the alignment identity using the rule: (a) if an exact match is found, a value of 0 will be reported; (b) if an insertion is found, a value of 1 will be reported; (c) if a deletion is found, a value of −1 will be reported; (d) if a mismatch is found, a value of 2 will be reported (Fig. 2).

### Neoantigen binding affinity by epitope prediction software

Numerous allele-specific epitope prediction software tools were published in characterizing the binding affinity of neoantigen binding to major histocompatibility complex (MHC). It’s not common to know HLA (human leukocyte antigen) class I haplotype of a patient, yet the epitope binding affinity prediction is achieved by using all known HLA in this study with software tool netMHC (Fig. 3).

**Fig. 3.**
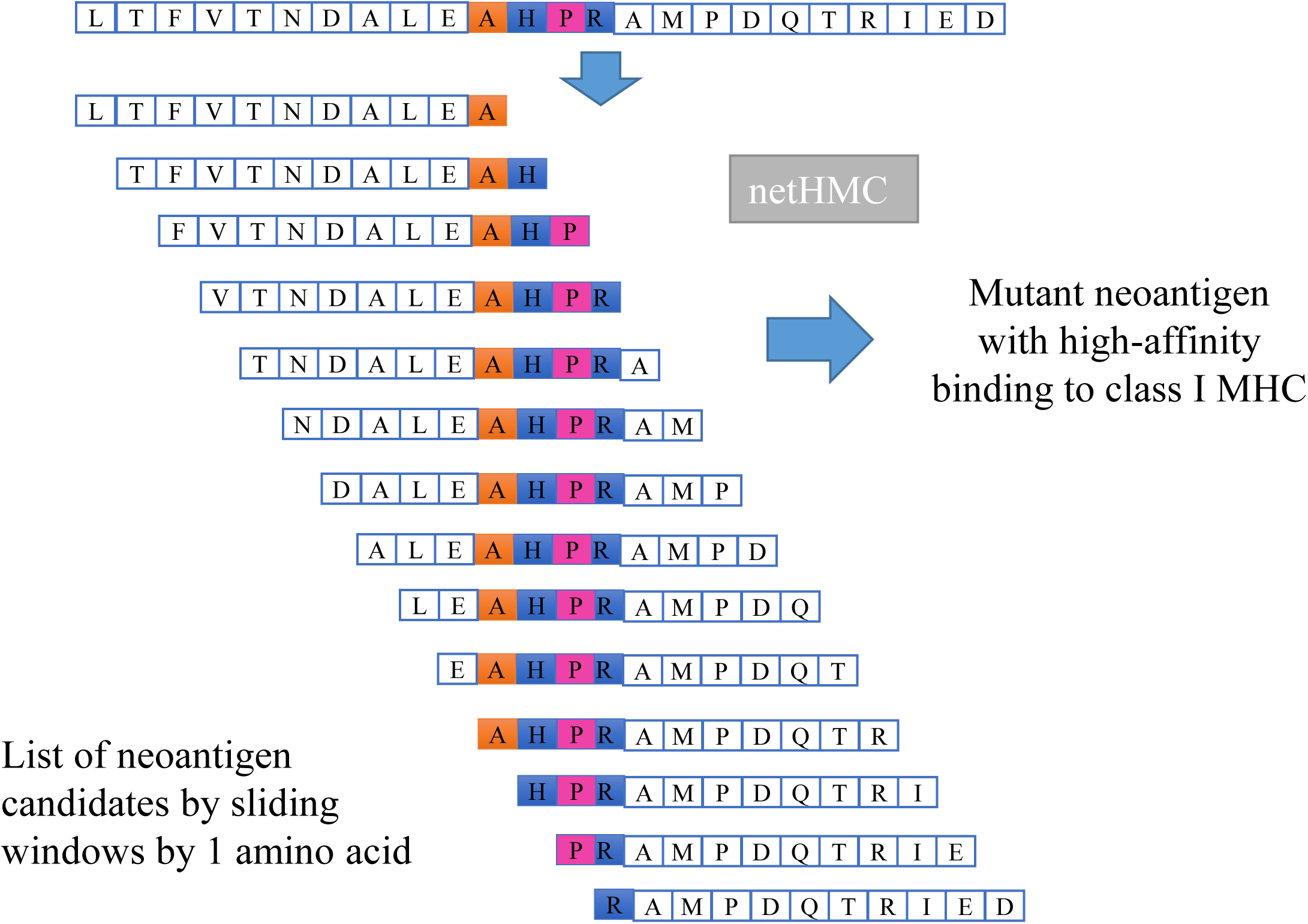
Generation of putative neoantigen precursor candidates from the peptides with sequence abnormalities: given a peptide with mismatched, insertions and deletions after protein sequence alignment, a list of peptides of length 11 amino acids were generated. Each of the neoantigen candidate peptide carries at least one amino acid from the alternative sequence (in color).

## Dataset

A dataset containing the full-length whole transcriptome from three diverse human tissues (brain and liver) was directly downloaded from the Pacific Biosciences (PacBio) official website (http://datasets.pacb.com.s3.amazonaws.com/2014/Iso-seq_Human_Tissues/list.html). This dataset is ideal for exploring differential alternative splicing events.

## Results

Fig. 4 indicated the flow of raw sequences read to neoantigens as the dataset run through our pipeline. Initially, the brain long-read sequencing data contained 10289 SMRT reads which are mapped to 7819 genes (Fig. 4). After protein sequence alignment with reference protein database such as the Uniprot Swiss-Prot reviewed proteins, which have been carefully curated and manually reviewed. We identified a total of 712 (553 unique) neoepitopes. As our method relied on a canonical protein sequence database for neoepitope discovery, some of our identified neoepitopes could be found in predicted, un-curated protein database. Users may address this shortcoming by using a comprehensive protein sequence database (a significant increase of running time will be expected), or performing the NCBI protein search to remove the false neoepitope predictions that exist in database of interest from predicted isoforms. Overall, in this dataset, we found that 283 (51.1%) of the predicted neoepitopes appearing in existing computational predictions or uncurated database other than Uniprot Swiss-Prot database. Users still have the flexibility to reconsider these neoepitopes since they are not shown in any protein sequences from reviewed and curated database such as swiss-uniprot. Nevertheless, after removing these 283 ‘neoepitopes’, 270 unannotated and novel neoepitopes were used as candidates for subsequent neoantigen identification. Among these peptides, more than half of them are translated from alternative splicing with exon skipping or alternative 5’/3’ sites. These 270 neoepitopes were uploaded to netMHC with default parameters. Overall, we identified 86 neoantigens with high binding affinity to existing HLA types, and 322 neoantigens with relatively weak binding affinity. In fact, multiple strong and/or weak binding neoantigens could be reported in a single neoepitope because each neoantigen is different from each other only by a single amino acid (Fig. 4).

**Fig. 4.**
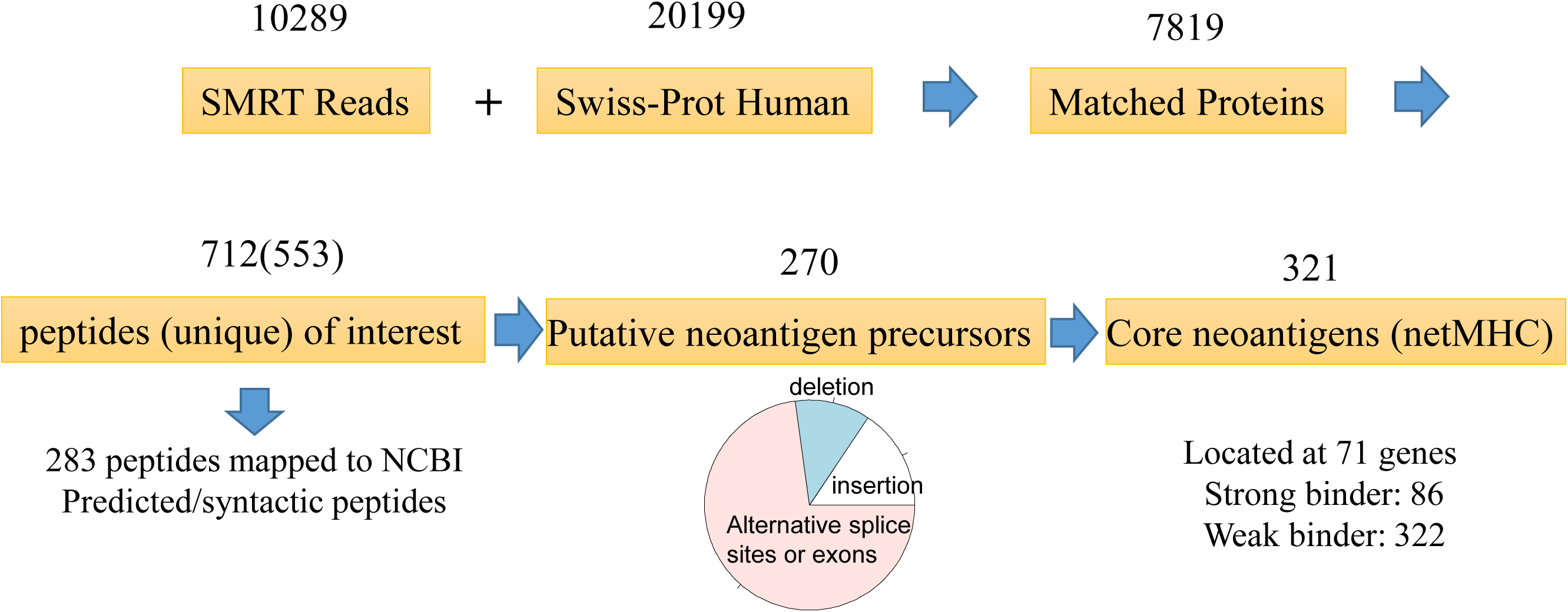
Flowchart of the identification of neoantigens using neoantigeR: using the PacBio’s public long read sequencing dataset generated from brain sample, we have shown that a set of 270 peptides are potential neoantigens out of the 10289 SMRT long reads. Binding affinity results from neoantigen prediction tool netMHC showed us that 321 candidate neoantigens have strong binding scores.

To give a systematic demonstration of the discovery of neoantigens from scratch, we showed an example from the initial mapping of SMRT read sequence to the identification of neoantigen on by gene PITPNM1. The gene prediction file indicated PITPNM1 has an alternative spliced event by skipping several exons when compared with the canonical gene (Fig. 5). In addition, the protein sequence search in the existing database didn’t find any exact match for the new isoform-specific protein sequence. The in-silico protein translation illustrated an in-frame deletion resulting in a significantly truncated protein (Fig. 6). The new truncated protein contained a new sequence fragment that was formed by joining the two portions of the protein sequence previously separated by the deleted sequence (Fig. 7). The neoepitope was subsequently sliced into a sliding window of 10 amino acid fragments and processed by netMHC for binding affinity prediction. Results showed a unique neoantigen candidate sequence with a strong binding affinity to HLA-A0201.

**Fig. 5.**
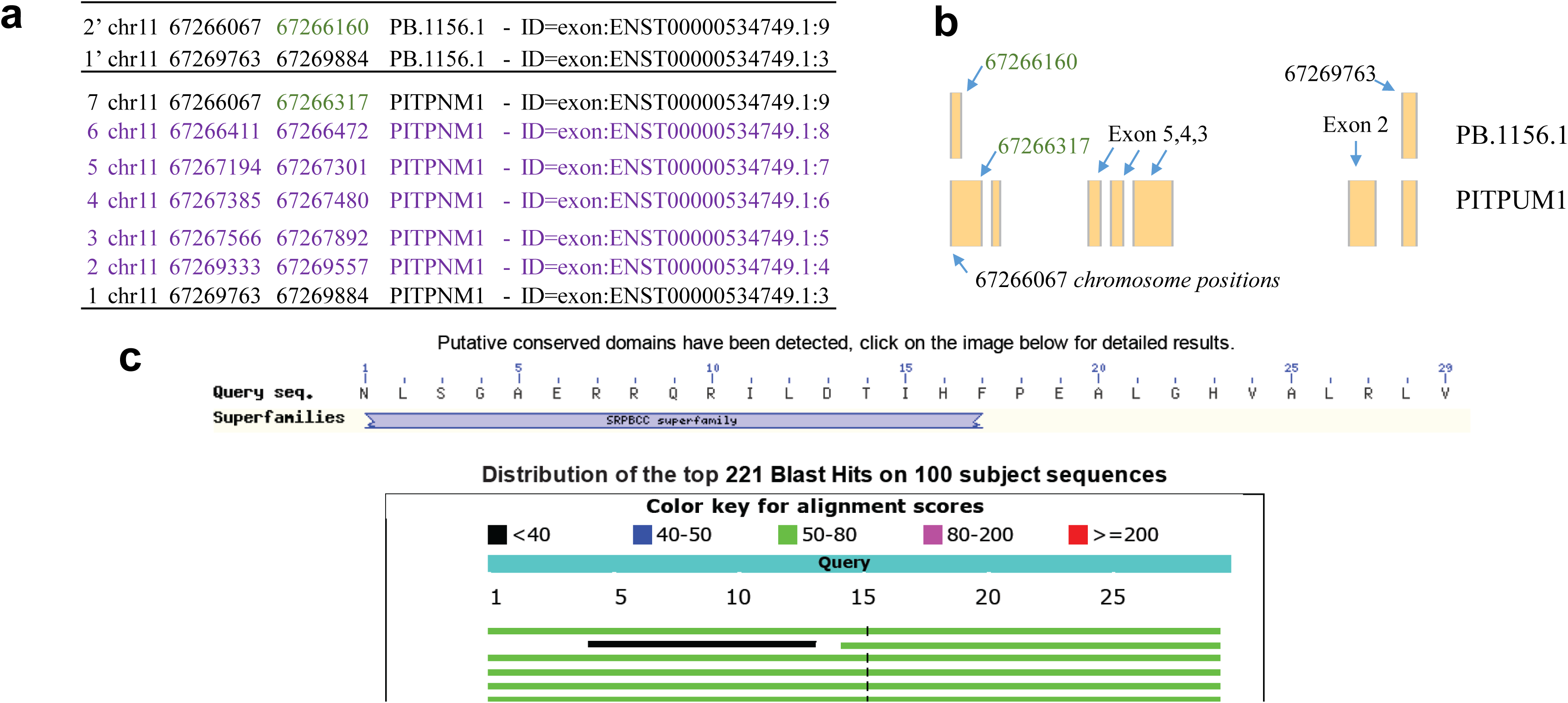
Exon skipping and alternative 3’ site resulting in a truncated protein in PITPUM1 gene. a. this table showed the exon coordinates of the predicted PacBio transcript and the matching protein PITPNM1 (green color indicates the alternative 3’ site and purple indicates the skipeed exons. b. illustration of the skipped and alternative spliced exons. c. NCBI BLASTP confirmed that there is a novel protein isoforms.

**Fig. 6.**
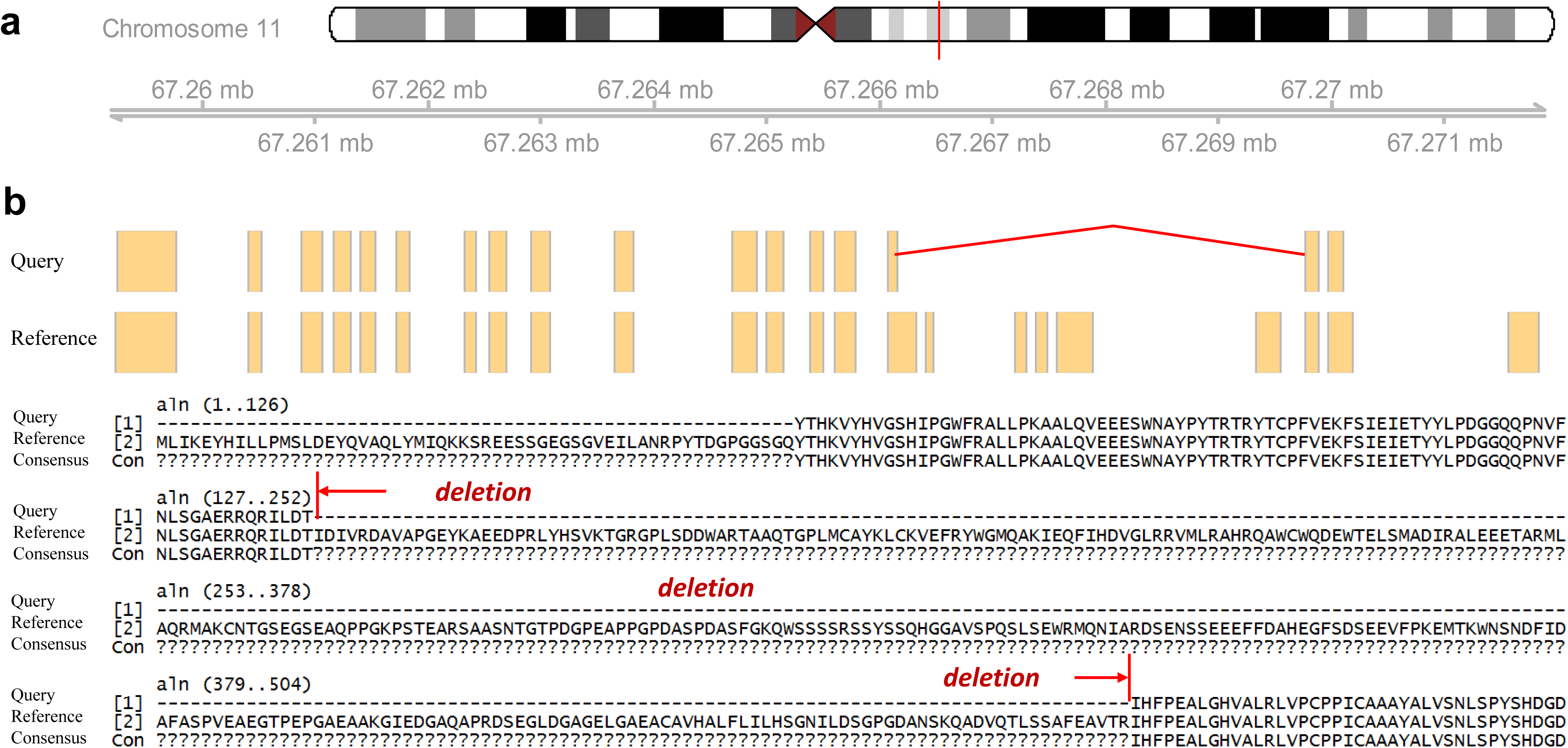
Exon skipping and alternative 3’ site resulting in a truncated protein PITPNM1. a.neoantigenR automatically produces the graphical gene tracks and the comparison of disease vs normal (or reference genes). b. the alternative 3’site and exon skipping generated an in-frame deletion.

**Fig. 7.**
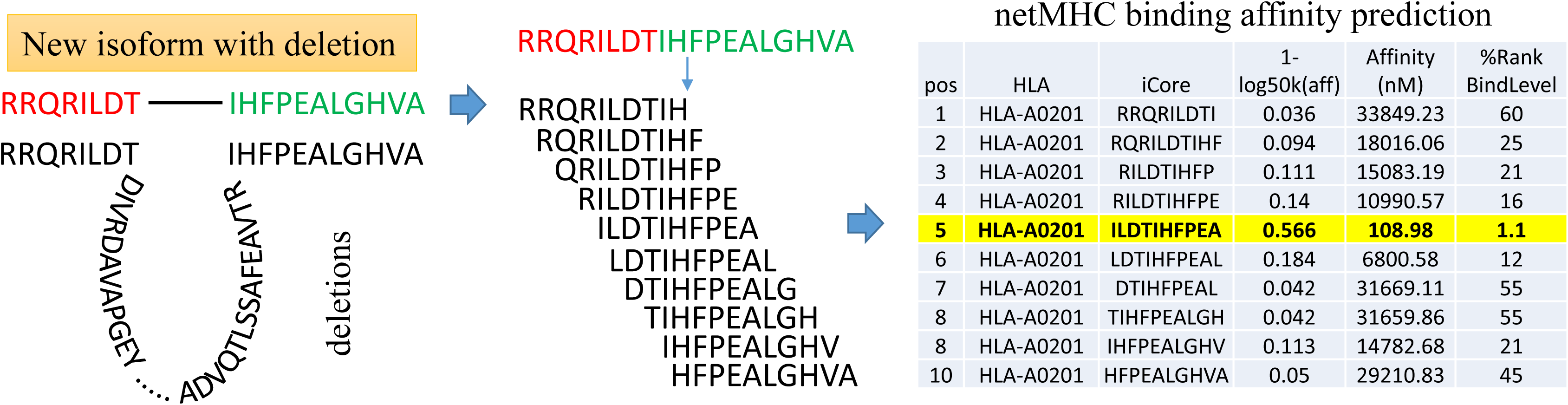
Detection of a putative neoantigen in the alternative isofroms of gene PITPUM1. The alternative truncated protein shown in Figure 6 lead to the generation of a new sequence fragment (the concatenation of flanking regions surrounding the deletion). Among the putative neoantigen candidates, one of them has a strong binding score with MHC indicating a large probability of being a neoantigen.

A similar analysis was performed on the liver sample (the input alignment file is about half the size of the brain sample) with a total of 336 (245 unique) sample-specific isoforms neoepitopes. Only 13 neoepitopes were shared between the liver sample and the brain sample, indicating that the alternative splicing is typically tissue-specific and is closely regulated by developmental stage and pathological conditions. The study of the tissue specific alternative splicing is thus important for the characterization of mutation burden and novel neoepitopes that may trigger adaptive immune response. As evidenced in table 1, there is a multitude of epitopes reported by neoantigenR. As mentioned earlier, for the demonstration of this workflow, structural changes such as insertion, deletion and amino acid changes resulting from only missense mutations were considered for final analysis. As the last part of our local pipeline, NetMHC v3.4 was used as the epitope prediction software to generate HLA class I restricted epitopes from our peptide epitope precursors. Using neoantigenR and netMHC (with default binding affinity cutoff), we were able to produce a greater list of high affinity HLA class I binding neoantigen candidates for future experimental validation. Essentially, there are 408 neoantigens reported by NetMHC v3.4 from 71 genes (see table 1).

**Table 1.**
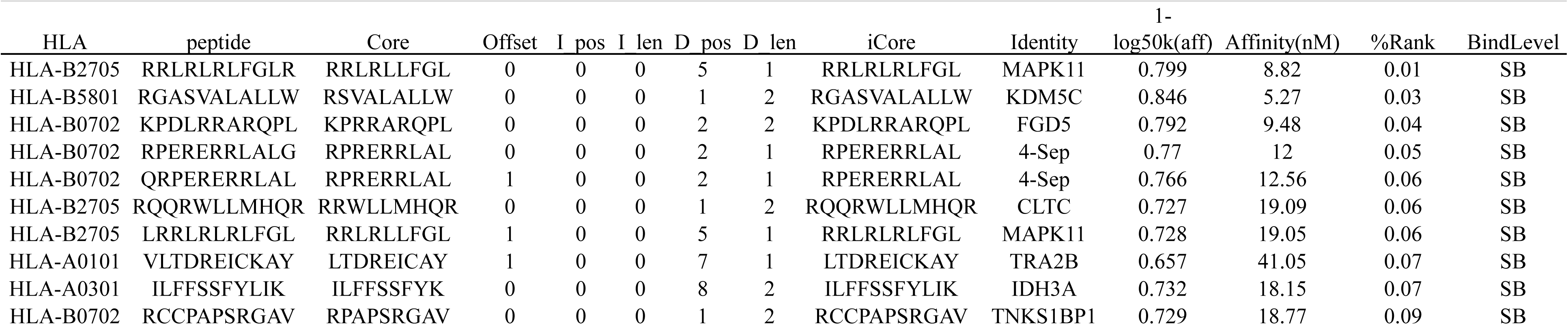
Top 10 highest affinity neoantigenes

Overall, neoantigenR used a sophisticated algorithm to retain high-quality sequence variations that would otherwise be challenging to produce peptide epitope precursors from the enormous searching domain of sequence anomalies using high-throughput sequencing data. By implementing the neoantigenR, we were able to rapidly assemble both genomic anomalies and single amino acid changes of potentially immunogenic neoepitopes within the landscape of all neoepitopes. We have demonstrated that it is possible to uncover essential splicing aberrance and genomic abnormalies by expanding the focus from somatic point mutations (pVAC-Seq) to the entire neoantigen landscape.

## Discussion

Our pipeline enables a one-stop prediction of disease-specific alternative spliced neoepitopes that could be used to directly screen for neoantigens in the future. This approach allows us to evaluate tumor-specific epitopes in a wide range of sequencing technologies such as short-read or long-read sequencing. High-throughput sequencing produces an enormous amount of DNA sequences that harbor point mutations, sequence insertions, fragment deletions, and alternative usage of exons (Hundal et al. 2016). It can be challenging to identify the best set of potential tumor-specific immunogenic neoantigens for cancer precision treatment and vaccine design. Altered proteins with unique amino acid sequences caused by changes to DNA may lead to various diseases including cancer. If properly presented by the immune system, these altered peptide sequences may elicit immune responses such as T-cell recognition, following protein degradation and major histocompatibility complex binding.

In the last few years, the development of high throughput sequencing technology coupled with computational data analysis enables an in-silico identification unique to tumor peptide sequences (“Neoantigen Discovery in Human Cancers : The Cancer Journal” 2017). The neoantigens detected in cancer when comparing with matched normal samples could serve as a precursor to designing a personalized cancer vaccine. The overall neoantigen load could also be used to indicate the biomarkers for cancer immunotherapy treatment. It has long been established that these neoantigens could be derived from transcripts’ high-throughput sequencing data. In RNA-Seq transcriptome data, it is common to observe thousands of alternative isoforms sequence that are novel transcribed regions or variants of existing genes, however, the majority of these isoforms will result in significantly truncated proteins or sequences interrupted by many stop codons when performing in-silico translation. It is essential to survey the fraction of these isoforms that will produce peptide sequences highly similar to reference proteins with meaningful sequence alternations, but it is a laborious process to screen and predict neoantigens from sequencing data. Overall, neoantigenR will assist in identifying antibodies for personalized high precision cancer vaccines.

